# Conserved task-specific profiles outweigh seasonal shifts in cuticular hydrocarbons of European honey bee subspecies

**DOI:** 10.64898/2026.05.05.722949

**Authors:** Daniel Sebastián Rodríguez-León, Aleksandar Uzunov, Cecilia Costa, Dylan Elen, Leonidas Charistos, Thomas Galea, Martin Gabel, Maria Alice Pinto, Ricarda Scheiner, Thomas Schmitt

**Author notes:** Corresponding author (DS RodrÍguez-León) | (T Schmitt).

## Abstract

Cuticular hydrocarbons (CHCs) are essential for insect waterproofing, yet how they change seasonally in social insects remains poorly understood. Due to its distinct seasonal worker phenotypes (summer and winter bees) and diverse subspecies, the western honey bee (*Apis mellifera*) is an ideal model to study seasonal CHC plasticity across populations with distinct local adaptations. We performed a common garden experiment to investigate the seasonal plasticity in CHC profiles across five European subspecies (*A. m. carnica, A. m. iberiensis, A. m. ligustica, A. m. macedonica, A. m. ruttneri*). We compared the CHC composition of workers performing tasks inside (“in-hive”) or outside (“out-hive”) the colony during summer and winter. Notably, out-hive workers consistently exhibited more waterproofing CHC profiles compared to in-hive workers, regardless of season or subspecies. The persistence of this stereotypical task-related differentiation in long-lived winter bees, which largely lack an age-based division of labor, indicates a robust, age-independent regulatory mechanism linked to the environment faced by the workers rather than a simple response to seasonal desiccation pressure. Moreover, we demonstrate CHC seasonal plasticity for the first time in honey bees. However, these seasonal shifts in hydrocarbon classes and chain length were not uniform; they varied across subspecies and critically depended on the task the workers performed.

## Introduction

Insects are highly sensitive to variation in temperature and humidity, being particularly vulnerable to water loss by transpiration through their cuticle, due to their high surface-to-volume ratio [1,2]. The transpiration is minimized by cuticular hydrocarbons (CHCs), which form a semi-liquid waterproof coating on the epicuticle of insects [1,3,4]. The effectiveness of this coating in preventing water loss under specific temperature and humidity conditions is determined by its composition [3–5]. The tighter the aggregation of its constituent hydrocarbons, the more waterproof the CHC layer is [6,7]. Compared to *n*-alkanes, hydrocarbons with unsaturations (double bonds), or methyl-branches, reduce hydrocarbon aggregation by limiting the formation of van der Waals bonds [5,7,8]. Molecule aggregation also increases with the hydrocarbon chain length [9]. As waterproofing requirements vary with temperature and humidity, CHC plasticity is crucial for insects to cope with desiccation stress across seasons.

The western honey bee (*Apis mellifera*) is an excellent model to investigate seasonal adaptive plasticity and its variation across populations with distinct local adaptations. It has evolved into 31 subspecies adapted to a wide diversity of environments throughout Africa and a large portion of Eurasia [10–14]. Their divergent evolutionary history has resulted in both genetic and phenotypic differences across *A. mellifera* subspecies [10,15]. The different subspecies have evolved local adaptive traits to survive the different climatic conditions within their native distribution ranges [14,16,17]. European *A. mellifera* subspecies, in particular, are adapted to strong seasonal variations in temperature and resource availability, representing a remarkable example of seasonal plasticity [18–21].

In subspecies native to the temperate regions of the northern hemisphere, the worker population can be divided into two seasonal groups along the year with markedly different life spans and behavior: short-lived summer bees and long-lived winter bees [22]. Summer bees have an individual life span of approximately 6 weeks and engage in different tasks during spring and summer, following an age-related division of labor [22,23]. Typically, the youngest workers care for the brood and queen (nurse bees), while the oldest workers collect pollen, nectar, water and propolis outside the colony (forager bees) [24]. In contrast, winter bees (also referred to as “*diutinus*” bees), typically emerge in autumn, have an individual life span of up to 6 months, engage extensively in thermoregulation during winter, and display a generalist task repertoire during the spring [22,23,25]. These two seasonal groups present diverse adaptations to ensure the survival of the colony under the different environmental conditions they face [25–27].

*A. mellifera* subspecies have evolved distinct CHC profiles, which exhibit both qualitative and quantitative compositional differences [28]. The CHC profiles of summer honey bee workers vary with subspecies and task performance [28–30]. Yet, the largest CHC composition variation in summer honey bee workers correlates with their task performance [28,30]. The CHC profiles of nurse bees are characterized by a higher abundance of unsaturated hydrocarbons and longer chain lengths compared to forager bees [28,29]. These differences seem to respond to the contrasting environmental conditions (e.g., temperature and humidity) that the nurse and forager bees face [28,29]. Nurse bees perform tasks inside the hive and thus face a more controlled environment compared to forager bees, which are exposed to a harsher environment while gathering resources outside the hive. Forager bees are therefore thought to have more waterproofing CHC profiles to better protect themselves against desiccation [28–30]. These task-related differences in CHC composition might also aid the division of labor by informing the task performance of a specific worker to its nestmates [31].

Winter bees mainly engage in thermoregulation over the winter and in raising the next generation of bees in spring [22]. They display a rather generalist task repertoire in the late winter/early spring [22,23,32]. At that point, winter bee division of labor is unlikely to respond to their age, as they are all much older than the oldest summer foragers. However, some winter bees tend to remain inside the nest and others forage, during the spring, when they are raising the next generation of workers [22,32].

In winter bees, CHC profiles have not yet been described. It is therefore unknown how they differ from those of summer bees and whether they differ between individuals at all, given the lack of clear task partitioning. However, it could be assumed that winter bees that spend time outside the hive as foragers might differ in their CHC profiles from winter bees that remain inside the hive, because the two groups are confronted with different weather conditions such as temperature and humidity. Alternatively, winter bees might show a rather uniform CHC profile. Their CHC composition might only be driven by the winter-specific environmental conditions. Whether seasonal plasticity in CHC profiles occurs across *A. mellifera* subspecies adapted to differing summer-winter regimes of desiccation pressure, is an intriguing question.

Here, we analyzed the CHC profiles of summer and winter worker bees of five European *A. mellifera* subspecies (*A. m. carnica, A. m. iberiensis, A. m. ligustica, A. m. macedonica*, and *A. m. ruttneri*), and the CHC composition between workers performing tasks inside of the hive (“in-hive”) and those outside of the hive (“out-hive”). We aimed to gain new insights into adaptation to seasonality across subspecies. Since winter survival is critical for honey bee colonies in temperate climates, identifying both subspecies-specific and season-specific CHC patterns may have important implications for breeding and conservation. We hypothesized (1) that CHC profiles are task-specific in both in-hive and out-hive workers; and (2) that seasonal environmental changes elicit composition changes in the CHC profile of honey bee workers, explained by the change in the desiccation pressure. Hence, we expected that in-hive and out-hive workers differ in CHC composition, with out-hive workers having more waterproofing CHC profiles in summer and winter. We also expected the CHC profiles of honey bee workers to differ between summer and winter across subspecies. Finally, because out-hive workers experience more pronounced seasonal variation in environmental conditions, we expected their CHC profiles to show greater seasonal divergence than those of in-hive workers.

## Materials and methods

We performed a common garden experiment by keeping queen-right colonies of five different honey bee subspecies (*A. m. carnica, A. m. iberiensis, A. m. ligustica, A. m. macedonica*, and *A. m. ruttneri*) in the departmental apiary of the University of Würzburg from April 2018 to March 2019. This ensured that the colonies experienced the same environmental conditions and thus the differences in their CHC composition would mainly be driven by their genetic differences [33]. The queens were reared and mated in the area of origin of the respective subspecies, and then transferred to the research apiary according to official EU regulations: *A. m. carnica* from Würzburg, Germany; *A. m. iberiensis* from Bragança, Portugal; *A. m. ligustica* from Reggio Emilia, Italy; *A. m. macedonica* from Nea Moudania, Greece; *A. m. ruttneri* from Malta, Malta. A total of 20 queens (four per subspecies) were introduced in queen-less *A. m. carnica* hives for the initial set-up. To avoid forager drifting, the hives were placed on hive stands in pairs of the same subspecies, facing the opposite direction, and the hive stands were grouped by subspecies, for details on colony set up see [33]. Furthermore, the hive boxes were sorted in different color combinations, helping the returning foragers to locate their own hive.

Five workers per task (in-hive and out-hive workers) were collected from two different hives of each subspecies in summer and winter, for a total of 40 worker bees per subspecies. Collected bees were immediately killed by immersing them in liquid nitrogen and stored in labeled Eppendorf tubes at - 20 °C until hexane extraction. Workers were collected from each colony after a minimum of two months after queen introduction, once the turnover in the *A. m. carnica* recipient hives was ensured. In summer, bees were collected between July and September 2018, between 7 AM and noon. The summer in-hive workers corresponded to nurse bees, which were identified as workers poking their heads into an open brood cell for at least 10 seconds, demonstrating brood care. Summer out-hive workers corresponded to returning forager bees, which were caught at the hive entrance and identified by their noticeable pollen loads. The winter bees were collected in February 2019. This is a time during which they were expected to show a generalist task repertoire. Bees were collected in days when the temperature was over 6 °C, between 12:00 (noon) and 1:00 PM, to minimize cold stress on the hives while opening them. The winter in-hive workers were collected from the center of the inner frames of the hive. No brood was observed during the winter samplings, making it impossible to identify nurse bees. The winter out-hive workers were caught while landing at the hive entrance. No bee returning to the hive was observed carrying pollen loads during the winter samplings, impeding the identification of forager bees.

### CHC composition analysis

The CHCs were extracted from each individual bee by immersing their whole body in hexane for 10 min in 1.5 mL glass vials. Subsequently, each extract was concentrated by evaporating the solvent under a gentle stream of CO_2_ until a total volume of 100 µL, and transferred into a 300 µL glass insert. The insert was placed in a 1.5 mL glass vial. A 1 µL aliquot of each extract was injected into an Agilent 7890 A Series Gas Chromatography System (GC) coupled to an Agilent 5975 C Mass Selective Detector (MS) (Agilent Technologies, Waldbronn, Germany). The GC was equipped with a J & W, DB-5 fused silica capillary column (30 m x 0.25 mm ID; df = 0.25 µm; J & W, Folsom, CA, USA). The GC temperature was programmed from 60 to 300 °C with a 5 °C/min heating rate and held for 10 min at 300 °C. Helium was used as carrier gas with a constant flow of 1 ml/min. The injection was carried out at 300 °C in the split-less mode for 1 min with an automatic injector. The electron impact mass spectra (EI-MS) were recorded at 70 eV and with a source temperature of 230 °C.

The chromatograms were analyzed using the data analysis software package “MSD ChemStation F.01.00.1903” for Windows (Agilent Technologies, Waldbronn, Germany). The area of each peak was determined by integration, and the initial threshold of the integration parameters was set on 15. Initial area reject was set on 1, initial peak width was set on 0.02, and shoulder detection was off. The compounds were identified by their retention indices and diagnostic ions of their mass spectra. Double bond positions of monounsaturated hydrocarbons were identified by dimethyl disulfide derivatization [34]. Compounds that represented less than 0.01% of the total ion count of a sample and compounds in less than 50% of the extracts in a group were excluded from the analysis to avoid concentration effects and to compare group-specific profiles. The abundances of the compounds were quantified as relative abundances based on the areas of their corresponding peaks. For each sample, the area of each peak was divided by the sum of the areas of all peaks included in that sample. The resulting proportion for each peak within a sample was then expressed as a percentage.

Additionally, we calculated the relative abundance of the different hydrocarbon classes (i.e. *n*-alkanes, alkenes, alkadienes, and methyl-branched alkanes), as well as the mean chain length, in the CHC profile of each bee. The relative abundances of the different hydrocarbon classes were calculated as the sum of the relative abundances of all the individual compounds of a corresponding class that were found in the CHC extract of a bee. The mean chain length corresponds to the weighted mean of the chain length of all the compounds found in the CHC profile of each bee, using the relative abundance of the compounds as their weights.

## Data analysis

The CHC composition similarity between seasons, tasks, and subspecies was visualized with a bi-dimensional non-metric multidimensional scaling (NMDS), using the Bray-Curtis dissimilarity index to estimate the interindividual chemical dissimilarity. The difference in the CHC composition between seasons, tasks, and subspecies was tested via a PERMANOVA. The PERMANOVA included the interactions of the terms (season, task, and subspecies) and had unrestricted permutations. Additionally, pairwise comparisons with false discovery rate (FDR) Benjamini & Hochberg adjustments were performed to test the differences between season and task pairs.

We compared the relative abundance of the different substance classes of hydrocarbons, and the mean chain length between seasons (summer vs. winter) and tasks (in-hive vs. out-hive). We fitted quantile regression models for the 50% quantile, considering season, task, subspecies, and their interactions as independent variables (predictors). We further performed pair-wise post-hoc contrasts to estimate the differences between seasons for each task and subspecies group (e.g., in-or out-hive workers of *A. m. carnica*), and between tasks for each season and subspecies group (e.g., *A. m. carnica* summer or winter bees). These correspond to the model estimated absolute differences between groups and their 95% confidence intervals.

The data analysis was performed using R v4.5.2 [35] and RStudio IDE v2026.1.0.392 [36]. Data wrangling and plotting operations were done using the packages tidyverse v2.0.0 [37], ggtext v0.1.2 [38], gghalves v0.1.4 [39], ragg v1.5.0 [40], and patchwork v1.3.2 [41]. The NMDS and PERMANOVA were performed using the package vegan v2.7.2 [42]. The quantile regression models were fit with the packages quantreg v6.1 [43]. The marginal means and post-hoc pair-wise contrasts were estimated using the package emmeans v2.0.1 [44].

## Results

The CHC composition of in-hive and out-hive workers differed among seasons and subspecies (Figure 1; Tables 1 and S1). Notably, the effect of task on the CHC composition of the bees was stronger than that of season and subspecies (Task - SES: 150.292; Subspecies - SES: 41.539; Season - SES: 33.214; Table 1). While the CHC profiles of both in-hive and out-hive workers mainly differed in the relative abundance of their constituent hydrocarbons, both contained compounds that were not present in the CHC profiles of the other (see Supplementary file S1). The CHC profiles of in-hive and out-hive workers differed in the abundance of all substance classes, but these differences varied among seasons and subspecies (Figures 2 and S1 - S4). For most subspecies, the in-hive workers consistently had a higher abundance of *n*-alkanes in their CHC profiles than out-hive workers across seasons (Figures 2A and S1). The only exceptions were *A. m. iberiensis* winter bees and *A. m. ruttneri* summer bees, as in both cases the abundance of *n*-alkanes did not differ between in-hive and out-hive workers (winter *A. m. iberiensis* - 95% CI: -20.283, 2.893; summer *A. m. ruttneri* - 95% CI: -30.254, 6.789). The abundance of alkenes (Figures 2B and S2), alkadienes (Figures 2C and S3), and methyl-branched alkanes (Figures 2D and S4) was consistently higher in the CHC profiles of the in-hive workers compared to out-hive workers in summer and winter across subspecies, with few exceptions. *A. m. ruttneri* summer in-hive and out-hive workers did not differ in their abundance of alkenes (95% CI: -12.917, 41.198), alkadienes (95% CI: - 1.558, 3.596), and methyl-branched alkanes (95% CI: -1.094, 6.383). In turn, the summer in-hive and out-hive workers of *A. m. iberiensis* did not differ in the abundance of alkadienes (95% CI: -0.253, 7.770) and methyl-branched alkanes (95% CI: -2.429, 7.870), while the summer *A. m. carnica* in-hive and out-hive workers did not differ in the abundance of methyl-branched alkanes (95% CI: -0.133, 1.936).

**Table 1:** One-way PERMANOVA contrasting the CHC composition between seasons, tasks, subspecies, and their interactions. Df - degrees of freedom; SS - sum of squares; R2 - R squared; F - F-value; SES - standard effect size. p-values < 0.05 are marked in bold.

|  | Df | SS | R2 | F | SES | p-value |
| --- | --- | --- | --- | --- | --- | --- |
| Season | 1 | 0.895 | 0.057 | 25.427 | 33.214 | **0.001** |
| Task | 1 | 3.988 | 0.253 | 113.313 | 150.292 | **0.001** |
| Subspecies | 4 | 2.283 | 0.145 | 16.215 | 41.539 | **0.001** |
| Season x Task | 1 | 0.327 | 0.021 | 9.284 | 11.791 | **0.001** |
| Season x Subspecies | 4 | 0.864 | 0.055 | 6.141 | 13.504 | **0.001** |
| Task x Subspecies | 4 | 0.612 | 0.039 | 4.345 | 8.682 | **0.001** |
| Season x Task x Subspecies | 4 | 0.490 | 0.031 | 3.483 | 6.366 | **0.001** |
| Residual | 180 | 6.335 | 0.401 |  |  |  |
| Total | 199 | 15.793 | 1.000 |  |  |  |

**Figure 1:**
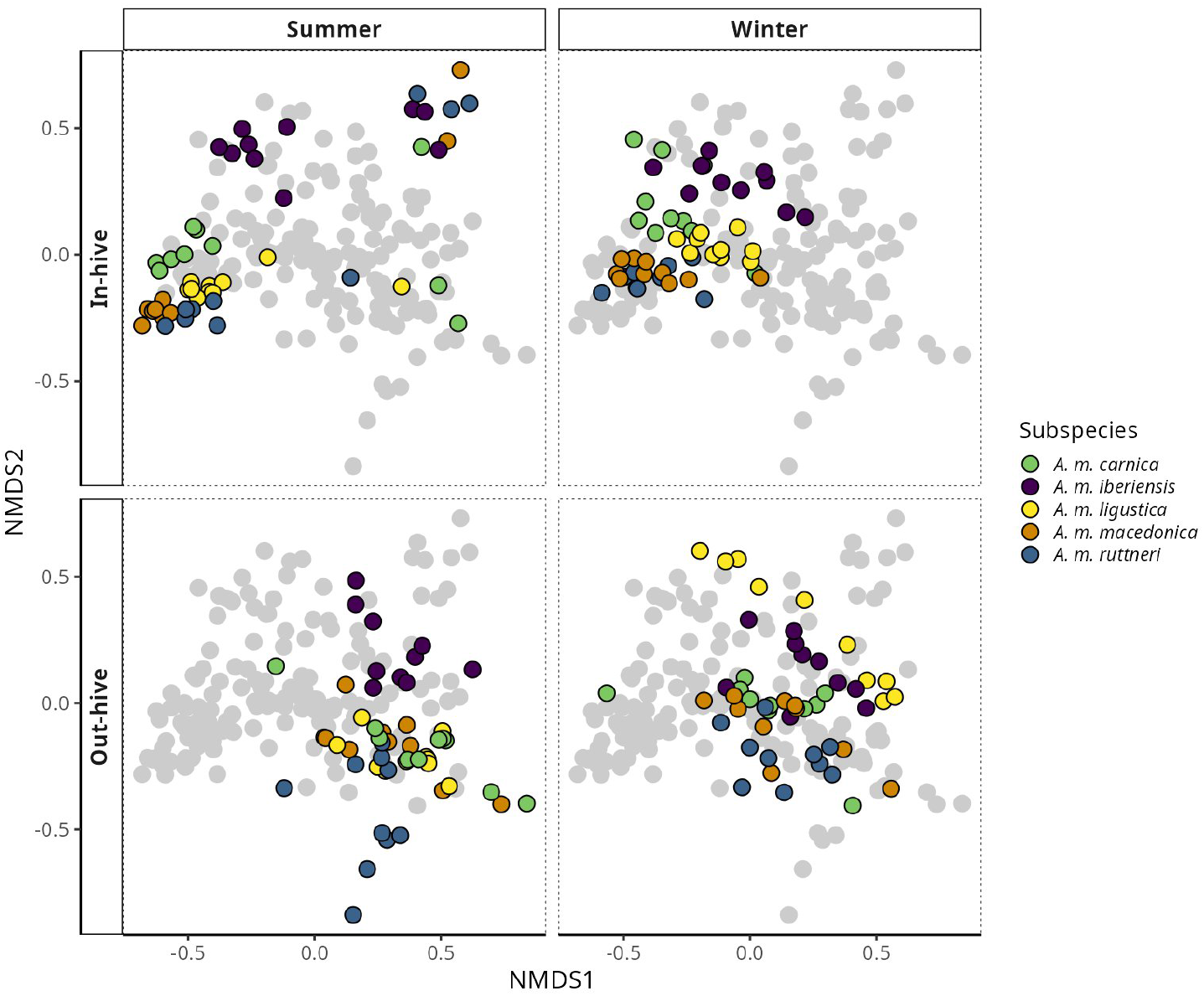
Bi-dimensional Non-metric Multidimensional Scaling (NMDS) analysis on a dissimilarity matrix of the CHC profile of honey bee workers (in-hive and out-hive workers) of five subspecies (*A. m. carnica, A. m. iberiensis, A. m. ligustica, A. m. macedonica*, and *A. m. ruttneri*) in summer and winter. Stress value: 0.140. The figure is organized as a 2X2 grid of panels. The columns represent seasons (left: summer; right: winter), and the rows represent tasks (top: In-hive workers; bottom: Out-hive workers). Each panel of the grid shows the same NMDS plot, but only the points representing bees of the corresponding season and task are colored, while the others are shown in grey.

**Figure 2:**
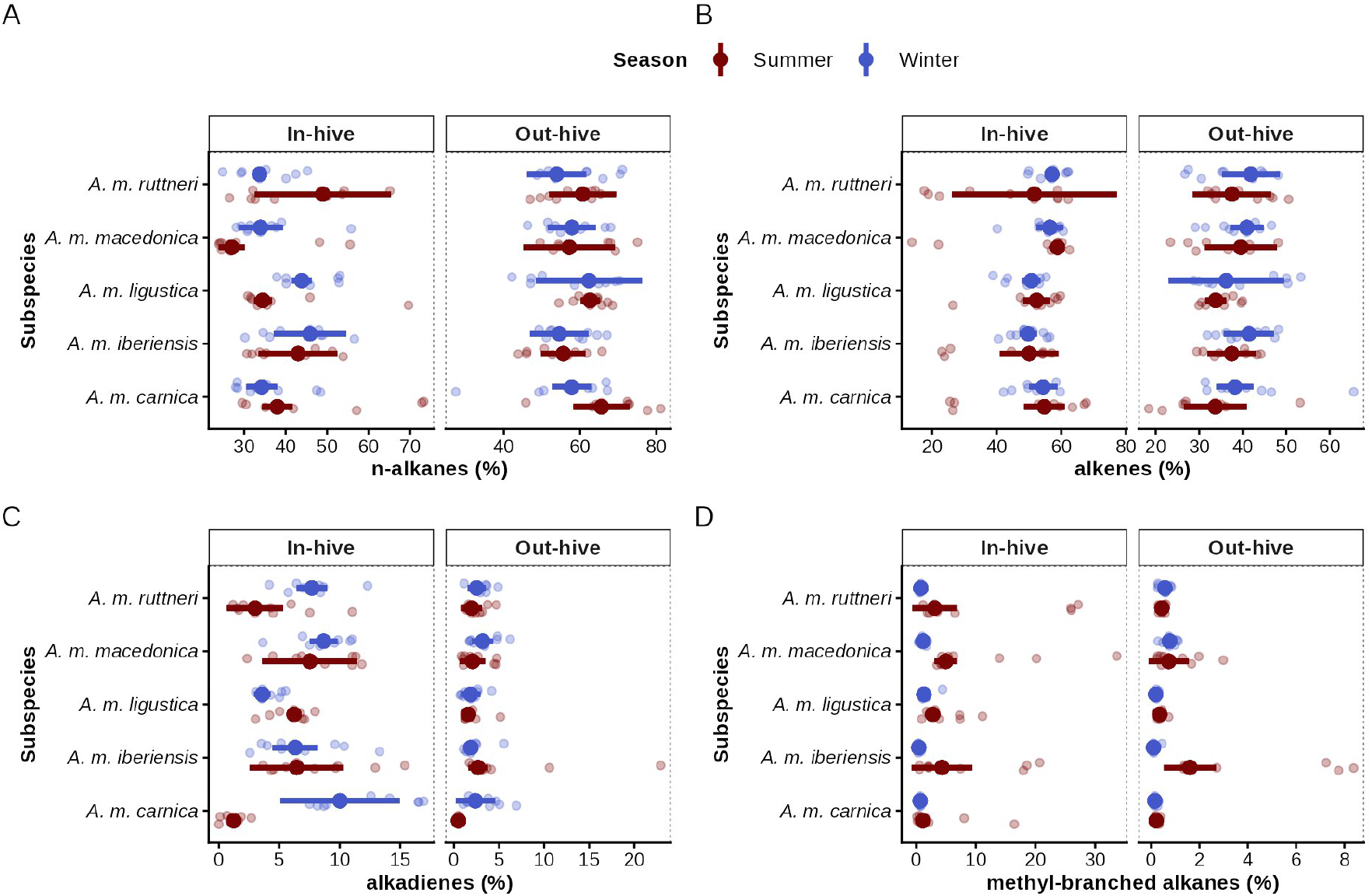
Abundance of hydrocarbon classes in the CHC profiles of honey bee workers. The figure is divided into four plots (**A, B, C**, and **D**), each depicting the results of a quantile regression analysis on the abundance of a hydrocarbon class in the CHC profile of summer and winter in- and out-hive workers of five *A. mellifera* subspecies. **A)** Relative abundance of *n*-alkanes. **B)** Relative abundance of alkenes. **C)** Relative abundance of alkadienes. **D)** Relative abundance of methyl-branched alkanes. The plots are divided into two sections regarding the task of the worker bees: in-hive workers on the left and out-hive workers on the right. Point-intervals represent the model prediction for the median and its 95% confidence interval. The raw data is visualized with transparent points, which correspond to the measured amount in the CHC profile of every bee.

The in-hive workers had longer hydrocarbons in their CHC profiles than the out-hive workers across subspecies, regardless of the season (Figures 3 and S5).

**Figure 3:**
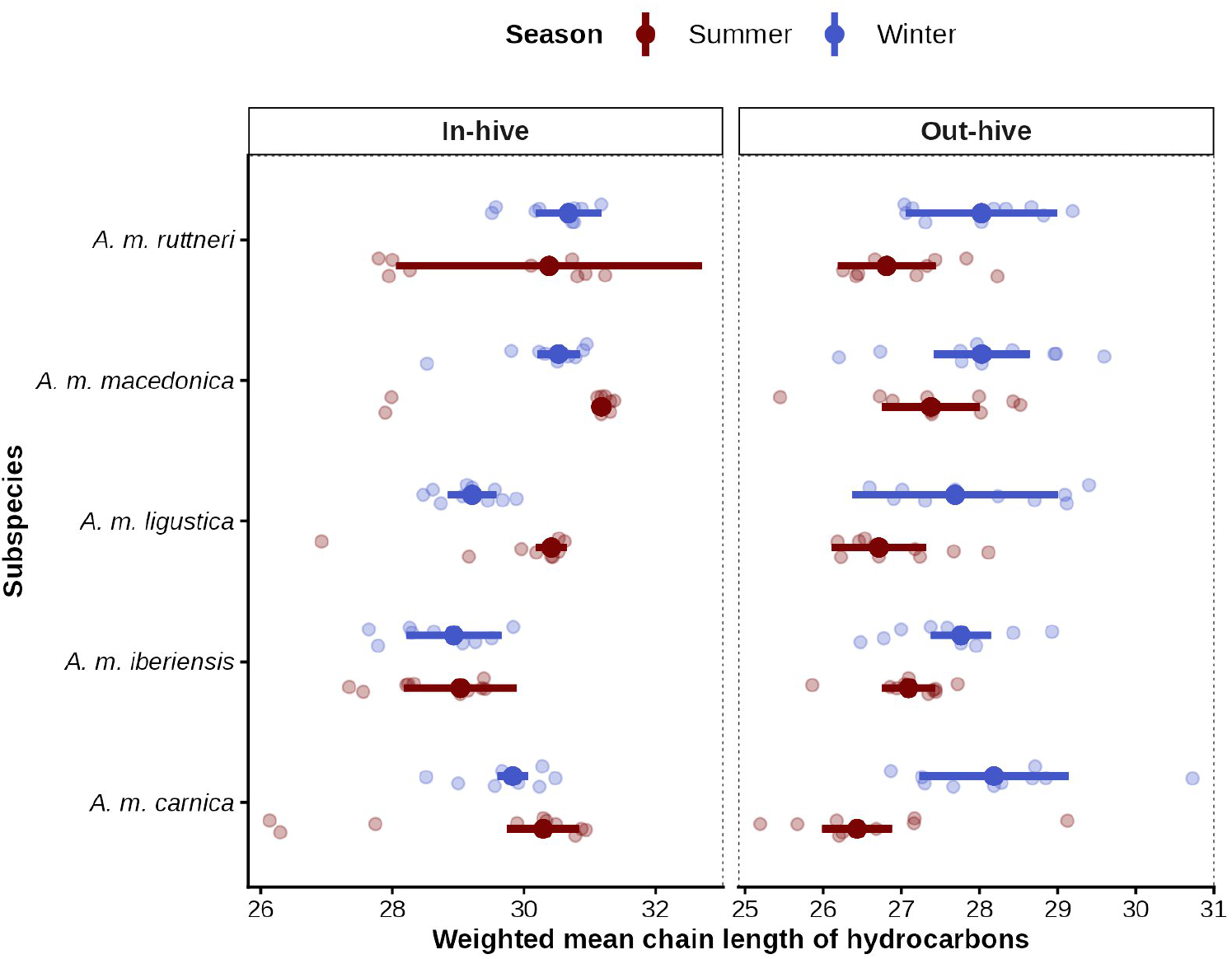
Mean hydrocarbon chain length in the CHC profiles of honey bee workers. The figure depicts the results of a quantile regression analysis on the weighted mean hydrocarbon chain length in the CHC profile of summer and winter in- and out-hive workers of five *A. mellifera* subspecies. The plot is divided into two sections regarding the task of the worker bees (in-hive workers on the left and out-hive workers on the right). Point-intervals represent the model prediction for the median and its 95% confidence interval. The raw data is visualized with transparent points, which correspond to the measured amount in the CHC profile of every bee.

Summer and winter bees of all five subspecies differed in their CHC composition (*p*-value<0.05; Figure 1; Tables 1 and S2). This differentiation was observed in both in-hive and out-hive workers of all five subspecies (Figure 1; Table S2). Although the CHC profiles of summer and winter bees mainly differed in the relative abundance of their compounds, each seasonal group displayed compounds that were not present in the other group (see Supplementary file S1). Seasonal differences in the relative abundance of *n*-alkanes, alkadienes, and methyl-branched alkanes occurred mainly in the in-hive workers and depended on the subspecies (Figures 2 and S1 - S4). The abundance of alkenes did not differ between seasons in both in-hive and out-hive workers across subspecies (Figures 2B and S2). The abundance of *n*-alkanes only differed between seasons for the in-hive workers of *A. m. ligustica* and *A. m. macedonica* (Figures 2A and S1). In both cases they were lower in summer bees than in winter bees (*A. m. ligustica* - 95% CI: -12.632, -6.191; *A. m. macedonica* - 95% CI: -13.114, -0.810). Alkadiene abundance differed between seasons in the in-hive workers, but not the out-hive workers, of *A. m. carnica, A. m. ligustica*, and *A. m. ruttneri* (Figures 2C and S3). The abundance of alkadienes was higher in summer than in winter in the CHC profiles of *A. m. ligustica* in-hive workers (95% CI: 1.781, 3.468), while the opposite was observed in the in-hive workers of both *A. m. carnica* (95% CI: -13.724, -3.877) and *A. m. ruttneri* (95% CI: -7.341, -2.083). In the case of the abundance of methyl-branched alkanes (Figures 2D and S4), it was higher in summer than in winter in the CHC profiles of *A. m. macedonica* in-hive workers (95% CI: 1.964, 5.671), *A. m. iberiensis* out-hive workers (95% CI: 0.433, 2.576), and both in-hive (95% CI: 0.200, 2.767) and out-hive workers (95% CI: 0.106, 0.228) of *A. m. ligustica*.

Summer and winter bees also differed in the chain length of the hydrocarbons in their CHC profiles, although this differentiation strongly depended on the task of the bees (Figures 3 and S5). In the case of the in-hive workers, the mean hydrocarbon chain length only differed between seasons for *A. m. ligustica* and *A. m. macedonica* (Figures 3 and S5), both having longer chain length hydrocarbons in summer than in winter (*A. m. ligustica* - 95% CI: 0.770, 1.632; *A. m. macedonica* - 95% CI: 0.320, 0.980). These two subspecies were also the only ones for which the mean hydrocarbon chain length did not differ between seasons for the out-hive workers (Figures 3 and S5). In all of the other three subspecies, out-hive worker CHCs had shorter mean hydrocarbon chain lengths in summer compared to winter (*A. m. carnica* - 95% CI: -2.802, -0.702; *A. m. iberiensis* - 95% CI: -1.181, -0.161; *A. m. ruttneri* - 95% CI: - 2.363, -0.0626).

## Discussion

In this study, we analyzed the CHC profiles of in-hive and out-hive honey bee workers in summer and winter in five different *A. mellifera* subspecies.

Notably, the CHC composition was remarkably more influenced by the task of the bees than by the season or the subspecies. In-hive workers exhibited hydrocarbons with longer chain lengths, a lower abundance of *n*-alkanes, and a higher abundance of alkenes, alkadienes, and methyl-branched alkanes compared to out-hive workers. These differences were consistent across seasons and subspecies, and suggest that out-hive workers have more waterproofing CHC profiles than in-hive workers, which should protect them better against desiccation [7,29,30,45]. These chemical differentiation patterns are very similar to those described for nurse and forager honey bees [28–30] and resemble those of ants workers inside and outside of the nest in several species [46–49]. Such differentiation likely reflects the higher exposure of out-hive workers to weather extremes (e.g., temperature and humidity) compared to in-hive workers [28,29,47]. Inside honey bee colonies, the temperature and humidity are under constant control to ensure the survival and proper development of the brood [50,51]. Out-hive workers are likely to experience more seasonally contrasting environments than their in-hive counterparts, in both summer and winter. Notably, the CHC composition of out-hive workers was more similar between seasons compared to that of in-hive workers. We suggest that this could derive from the difference in the division of labor between summer and winter bees. While summer bees show an age-related task specialization, winter bees mainly engage in thermoregulation during the winter and exhibit a generalist task repertoire in the late winter/early spring, when they raise the next generation of summer bees [22,23,25]. The stereotypical task-related CHC composition of honey bee workers is thought to aid the division of labor by informing the task performance of a specific worker to its nestmates [28–30]. Hence, the larger difference in CHC composition between summer and winter in-hive workers compared to out-hive workers might respond to the lower task differentiation of winter bees. In this sense, the summer in-hive workers would display a nurse-specific CHC profile, while the winter counterparts have CHC compositions that mainly respond to the desiccation pressure they face within the hive. In addition, the less contrasting CHC composition of summer and winter out-hive workers might respond to the higher desiccation pressure they face compared to in-hive workers, across seasons. The variability in their CHC profiles might be restricted because they are adapted to withstand a harsher environment outside of the hive, regardless of the season.

The observed differences between the CHC profiles of in-hive and out-hive workers were stereotypic across subspecies, suggesting that they respond to a conserved regulatory mechanism associated with the task transition of workers. *A. mellifera* task-specific CHC profiles have been linked to the variation in the expression of specific desaturase- and elongase-encoding genes [30], supporting the hypothesis of a regulatory mechanism yet unknown. As division of labor typically follows an age-dependent pattern in honey bees, this regulatory mechanism could be developmental [52]. However, *A. mellifera* division of labor flexibly responds to the needs of the colony, and individual bees transition from one task to another at different ages [24,53–55]. Under certain conditions, forager bees can even revert to nurse bees [24,56]. In this study, winter bees offer the opportunity to disentangle age and task in honey bee workers, as they were all much older than the oldest summer forager bees. Yet, we found that the differences in the CHC profiles of in-hive and out-hive workers followed a consistent pattern between seasons across subspecies. Therefore, these results suggest that the task-related CHC profiles respond to a non-age-dependent regulatory mechanism that allows honey bee workers to endure the desiccation pressure they face within their task-specific environment (i.e. inside or outside of the hive).

We provide for the first time evidence of seasonal plasticity in the CHC profiles of honey bee workers, as CHC composition differed between summer and winter bees, regardless of subspecies or tasks. We hypothesized that the observed seasonal CHC plasticity responds to the seasonal variation in desiccation pressure (temperature and humidity). Therefore the CHC profiles of summer and winter bees should differ in their waterproofing capabilities. Insects typically have a higher abundance of *n*-alkanes, a lower abundance of unsaturated and methyl-branched hydrocarbons, and/or longer hydrocarbons in their CHC profiles, when facing desiccation stress (low humidity or high temperature) [3,57–59]. Some *Drosophila* species, for example, increase the production of the longer hydrocarbons in their CHC profiles during winter [60,61]. However, the seasonal differences in the abundance of *n*-alkanes, alkadienes, and methyl-branched alkanes, as well as in the hydrocarbon chain length, largely depended on the subspecies of the bees, despite the common location, and whether they performed in-hive or out-hive tasks. Furthermore, the abundance of alkenes did not differ between seasons, in any of the subspecies. These results suggest that the different subspecies has evolved their own CHC composition response to seasonal fluctuations in desiccation pressure. Alternatively, they also represent evidence that the CHC composition shift between summer and winter bees does not predominantly respond to a differential desiccation pressure between seasons. Winter bees are physiologically different from summer bees. For instance, they present lower levels of juvenile hormone (JH) [18], higher levels of vitellogenin [25], and enlarged fat bodies [18,27], which increase their life span and tolerance to stress and starvation [25–27]. The seasonal shifts in CHC composition of honey bee workers may be influenced by physiological differences between summer and winter bees, or by the regulatory mechanism to which these physiological differences respond. Titers of JH, for example, are strongly correlated with the task transition from nursing to foraging in honey bee workers [62], which strongly correlates with the major variation in their CHC profiles [28–30,52]. In the African ant *Myrmicaria eumenoides*, the exogenous application of JH and its analogue methoprene induce an accelerated development of forager-like CHC profiles on the workers [48], and we hypothesize a similar mechanism for honey bees, which is currently under investigation. Hence, the difference in CHC composition between summer and winter bees might respond to the mechanism regulating the differentiation between these two seasonal worker groups.

## Conclusions

Here, we show that the CHC composition of honey bee workers is strongly influenced by the task performance inside or outside the hive, regardless of season. Across five European subspecies, in-hive workers consistently had profiles with higher abundance of unsaturated and methyl-branched hydrocarbons, and longer-chain hydrocarbons, whereas out-hive workers displayed profiles with higher abundance of *n*-alkanes. This stereotypical task-related pattern persisted in long-lived winter bees, indicating a conserved, age-independent regulatory mechanism. Moreover, the CHC profiles of out-hive workers varied less between seasons compared to in-hive workers, suggesting that they are adapted to withstand a harsher environment outside of the hive, regardless of the season. In addition, we show that honey bees have seasonal plasticity in their CHC profiles in five European subspecies, although the specific compositional shifts were subspecies-specific, regardless of the common location, and mainly occurred in the in-hive workers. This represents evidence that the different subspecies might have evolved their own CHC response to seasonal temperature and humidity fluctuations. Alternatively, we propose that the seasonal CHC plasticity is influenced by the physiological differences between summer and winter bees, or by the same mechanism regulating the transition of summer bees to winter bees rather than by desiccation pressure.

## Data availability

The data and code used for this study are not included in this manuscript, but will be made publicly available as supplementary material for the corresponding paper at the time of publication.

## Supplementary material

**Table S1:**
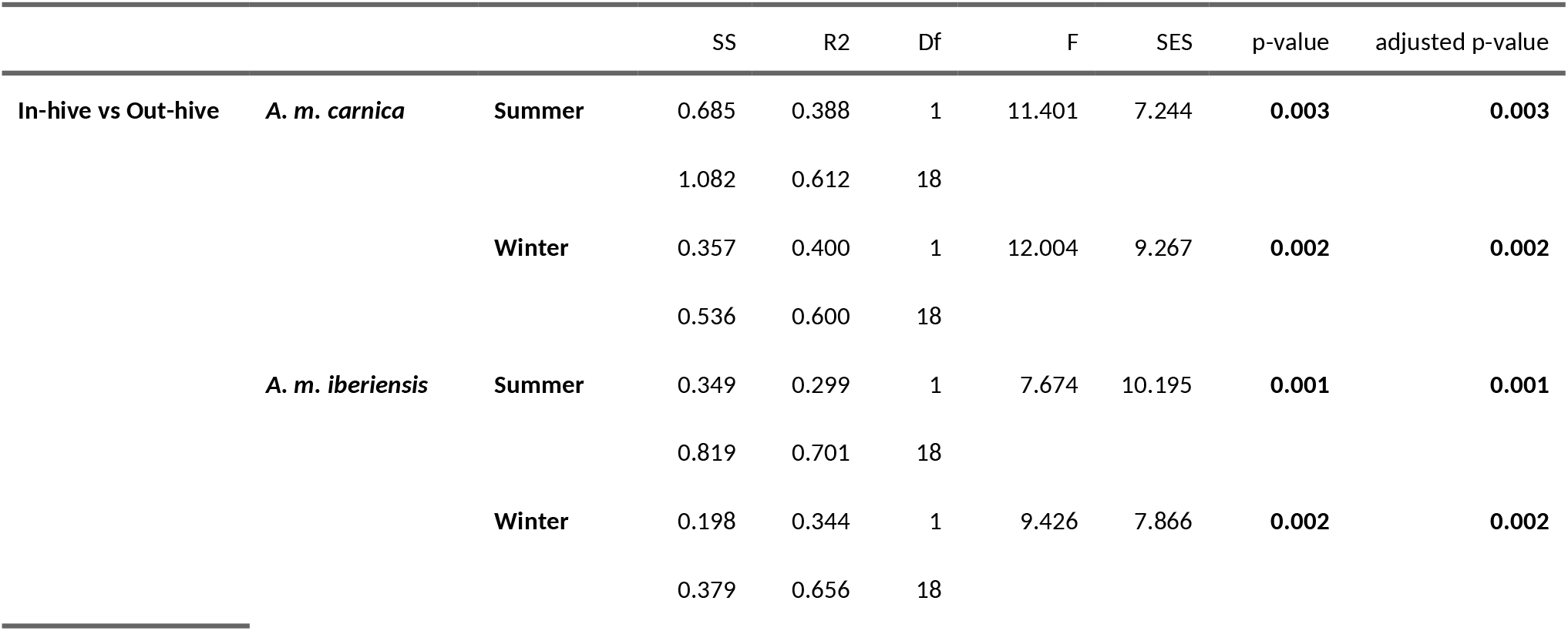
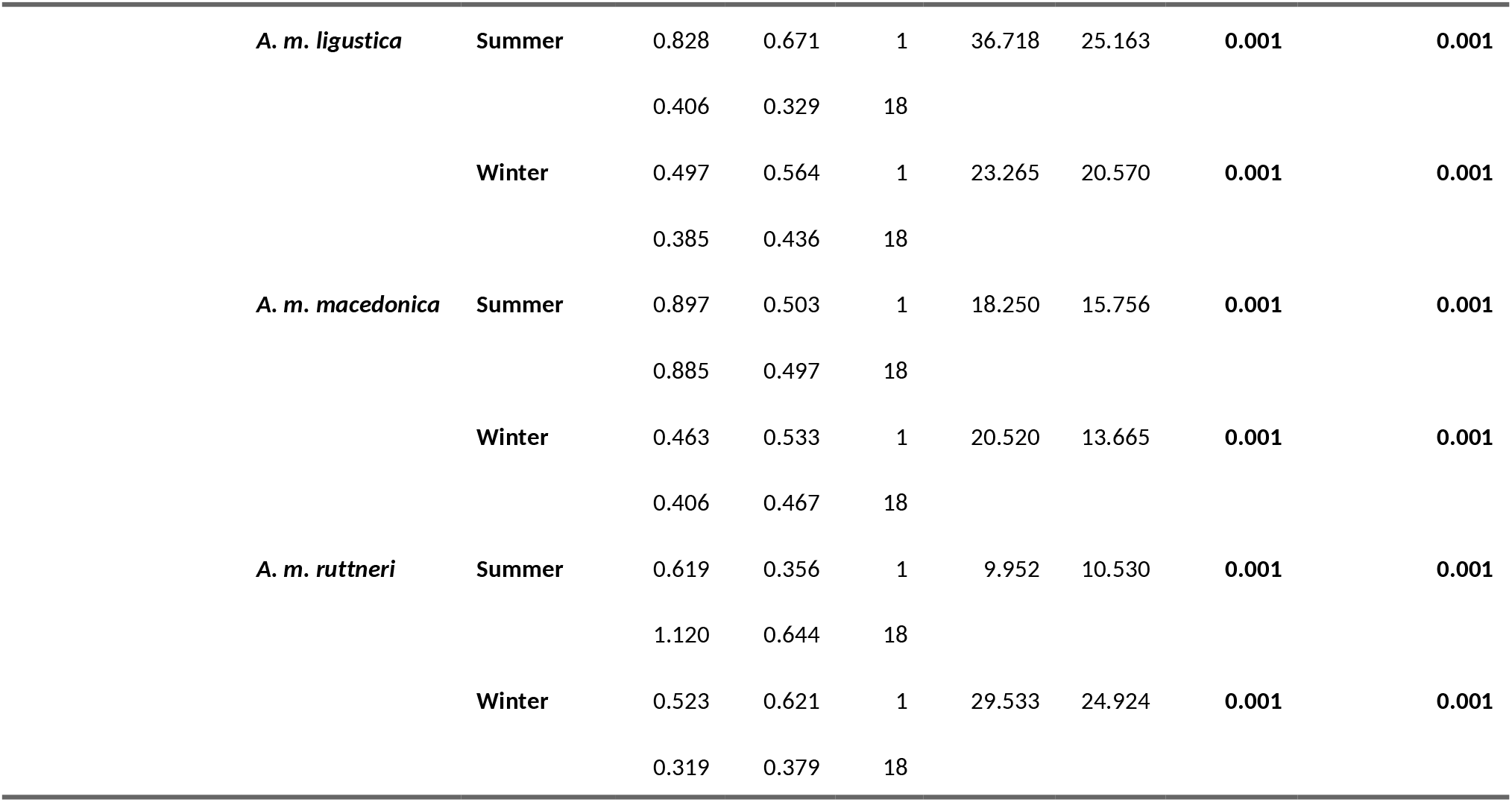
Pairwise PERMANOVA results contrasting the CHC composition between in- and out-hive worker bees for each subspecies and season. SS - sum of squares; R2 - R squared; Df - degress of freedom; F - F-value, SES - standard effect size. In the case of the SS and R2, the value for each comparison is displayed over that of the corresponding residuals. The adjusted p-values result from a FDR Benjamini & Hochberg adjustment. p-values < 0.05 are marked in bold.

**Table S2:** Pairwise PERMANOVA results contrasting the CHC composition between seasons for the in- and out-hive worker bees of each subspecies. SS - sum of squares; R2 - R squared; Df - degress of freedom; F - F-value, SES - standard effect size. In the case of the SS and R2, the value for each comparison is displayed over that of the corresponding residuals. The adjusted p-values result from a FDR Benjamini & Hochberg adjustment. p-values < 0.05 are marked in bold.

|  |  |  | SS | R2 | Df | F | SES | p-value | adjusted p-value |
| --- | --- | --- | --- | --- | --- | --- | --- | --- | --- |
| Summer vs Winter | In-hive | *A. m. carnica* | 0.316 | 0.243 | 1 | 5.768 | 6.154 | **0.001** | **0.002** |
|  |  |  | 0.985 | 0.757 | 18 |  |  |  |  |
|  |  | *A. m. iberiensis* | 0.192 | 0.208 | 1 | 4.738 | 4.736 | **0.002** | **0.003** |
|  |  |  | 0.730 | 0.792 | 18 |  |  |  |  |
|  |  | *A. m. ligustica* | 0.184 | 0.330 | 1 | 8.846 | 9.128 | **0.001** | **0.002** |
|  |  |  | 0.375 | 0.670 | 18 |  |  |  |  |
|  |  | *A. m. macedonica* | 0.195 | 0.212 | 1 | 4.843 | 4.221 | **0.001** | **0.002** |
|  |  |  | 0.724 | 0.788 | 18 |  |  |  |  |
|  |  | *A. m. ruttneri* | 0.242 | 0.225 | 1 | 5.217 | 3.863 | **0.006** | **0.007** |
|  |  |  | 0.835 | 0.775 | 18 |  |  |  |  |
|  | Out-hive | *A. m. carnica* | 0.262 | 0.293 | 1 | 7.445 | 4.800 | **0.006** | **0.007** |
|  |  |  | 0.633 | 0.707 | 18 |  |  |  |  |

|  | SS | R2 | Df | F | SES | p-value | adjusted p-value |
| --- | --- | --- | --- | --- | --- | --- | --- |
| ***A. m. iberiensis*** | 0.118 | 0.202 | 1 | 4.552 | 6.916 | **0.001** | **0.002** |
|  | 0.467 | 0.798 | 18 |  |  |  |  |
| ***A. m. ligustica*** | 0.656 | 0.612 | 1 | 28.407 | 23.062 | **0.001** | **0.002** |
|  | 0.416 | 0.388 | 18 |  |  |  |  |
| ***A. m. macedonica*** | 0.136 | 0.193 | 1 | 4.304 | 3.756 | **0.015** | **0.015** |
|  | 0.567 | 0.807 | 18 |  |  |  |  |
| ***A. m. ruttneri*** | 0.276 | 0.314 | 1 | 8.238 | 10.878 | **0.001** | **0.002** |
|  | 0.603 | 0.686 | 18 |  |  |  |  |

**Figure S1:**
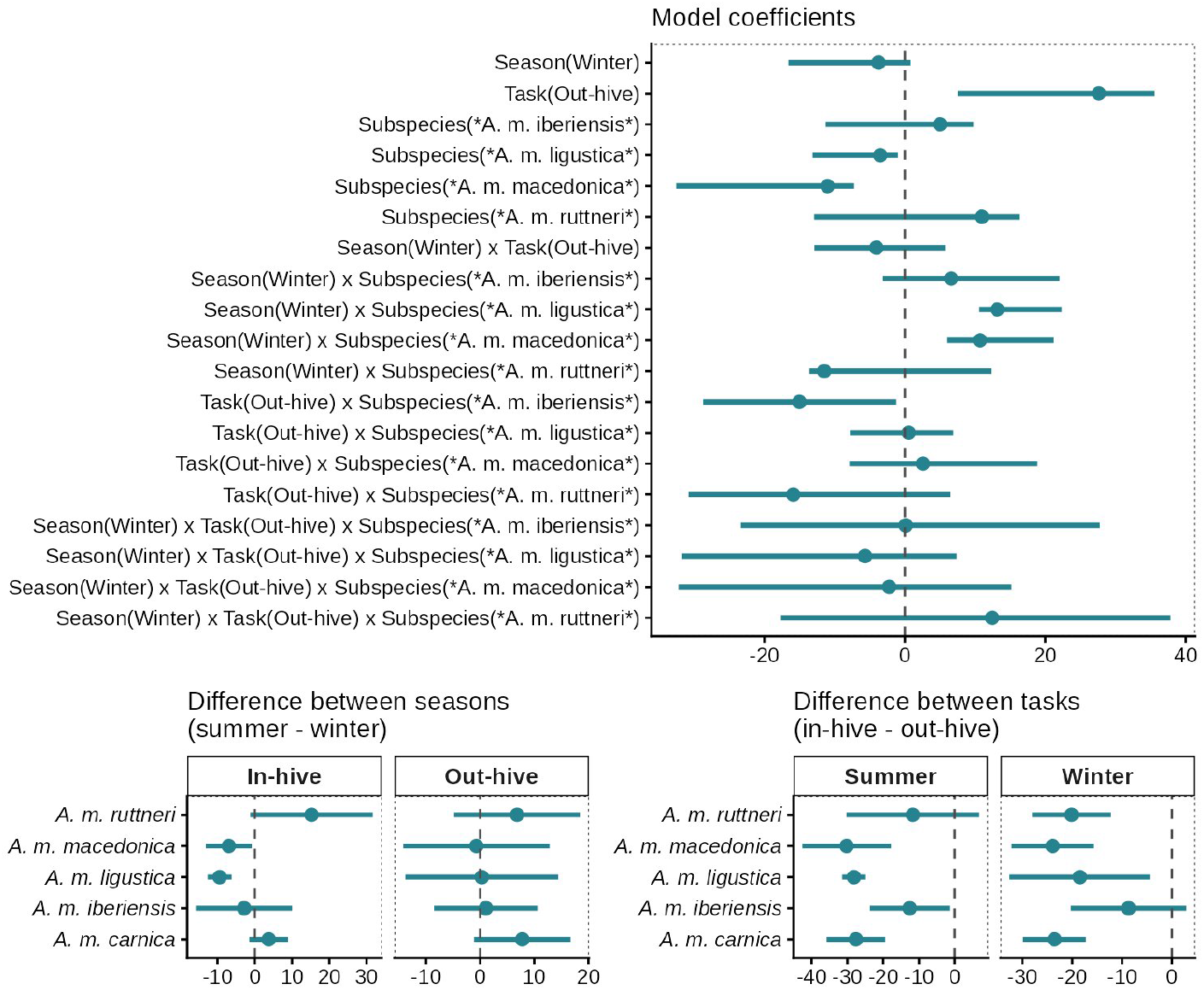
Differences in abundance of *n*-alkanes in the CHC profiles of summer and winter in- and out-hive workers of five *A. mellifera* subspecies. The figure is divided into three plots depicting the quantile regression model coefficients (top), and the pair-wise contrasts between seasons (bottom left) and tasks (bottom right). Point intervals represent the model estimates and their 95% confidence intervals. The null values are visualized with vertical dashed grey lines.

**Figure S2:**
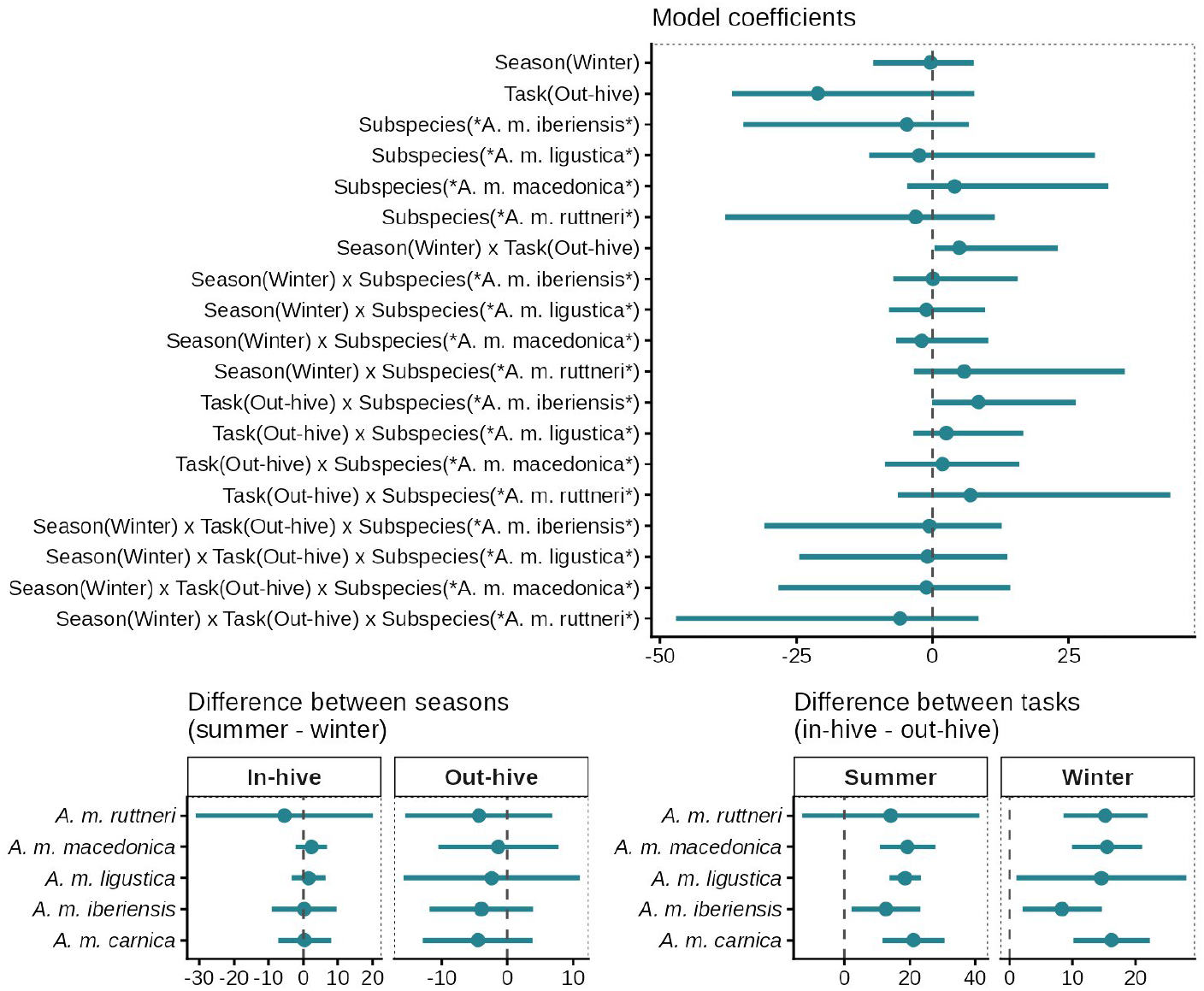
Differences in abundance of alkenes in the CHC profiles of summer and winter in- and out-hive workers of five *A. mellifera* subspecies. The figure is divided into three plots depicting the quantile regression model coefficients (top), and the pair-wise contrasts between seasons (bottom left) and tasks (bottom right). Point intervals represent the model estimates and their 95% confidence intervals. The null values are visualized with vertical dashed grey lines.

**Figure S3:**
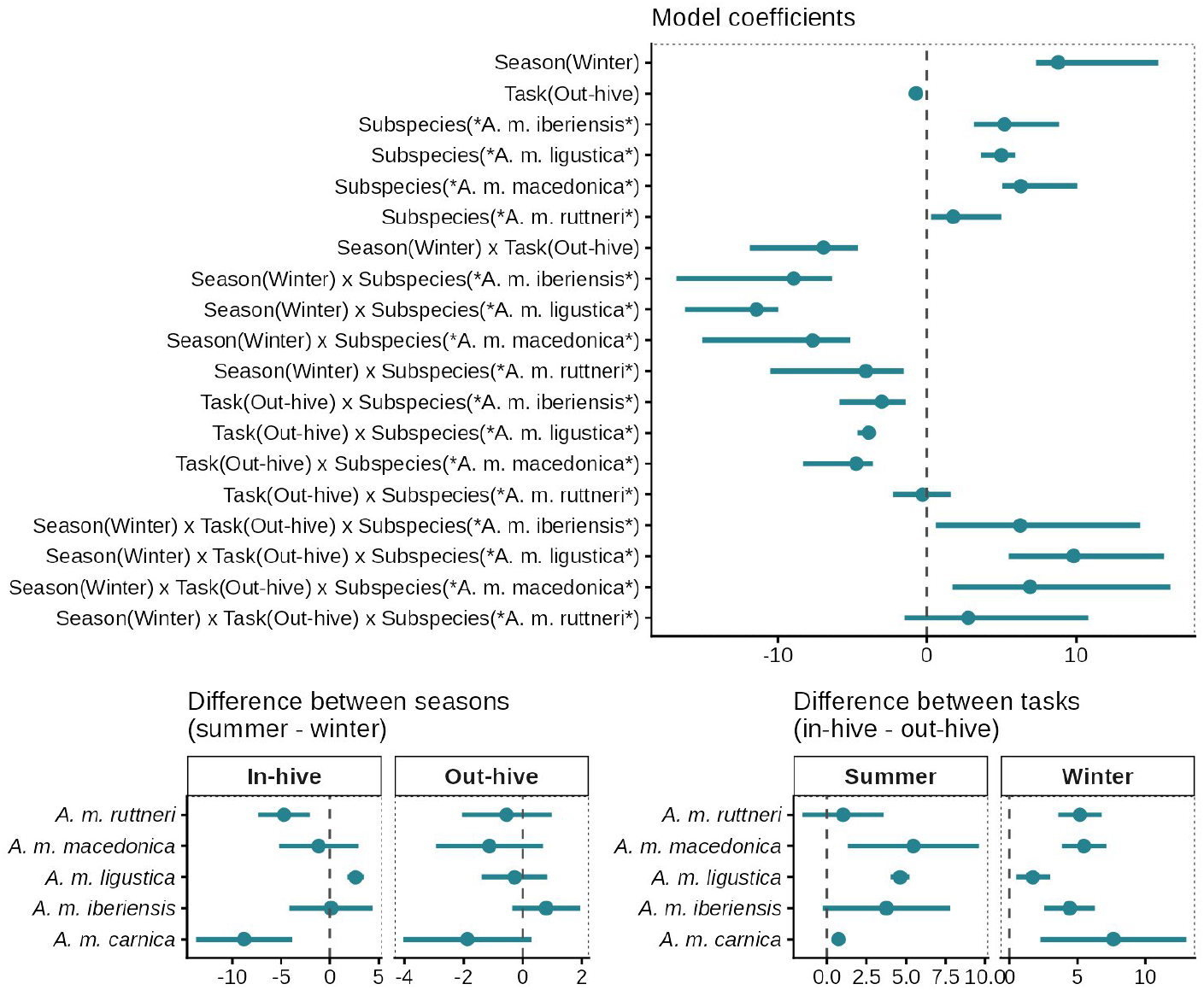
Differences in abundance of alkadienes in the CHC profiles of summer and winter in- and out-hive workers of five *A. mellifera* subspecies. The figure is divided into three plots depicting the quantile regression model coefficients (top), and the pair-wise contrasts between seasons (bottom left) and tasks (bottom right). Point intervals represent the model estimates and their 95% confidence intervals. The null values are visualized with vertical dashed grey lines.

**Figure S4:**
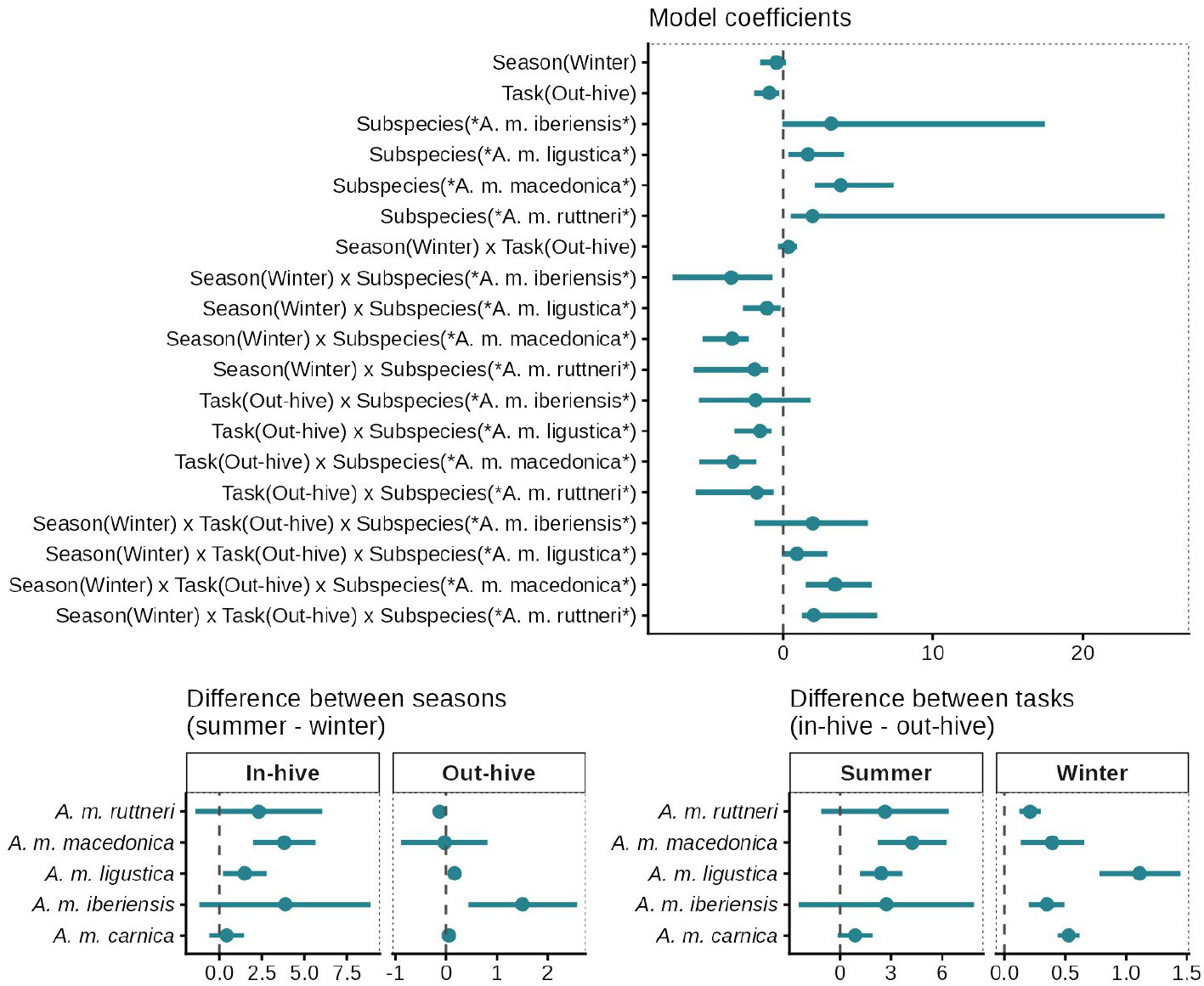
Differences in abundance of methyl-branched alkanes in the CHC profiles of summer and winter in- and out-hive workers of five *A. mellifera* subspecies. The figure is divided into three plots depicting the quantile regression model coefficients (top), and the pair-wise contrasts between seasons (bottom left) and tasks (bottom right). Point intervals represent the model estimates and their 95% confidence intervals. The null values are visualized with vertical dashed grey lines.

**Figure S5:**
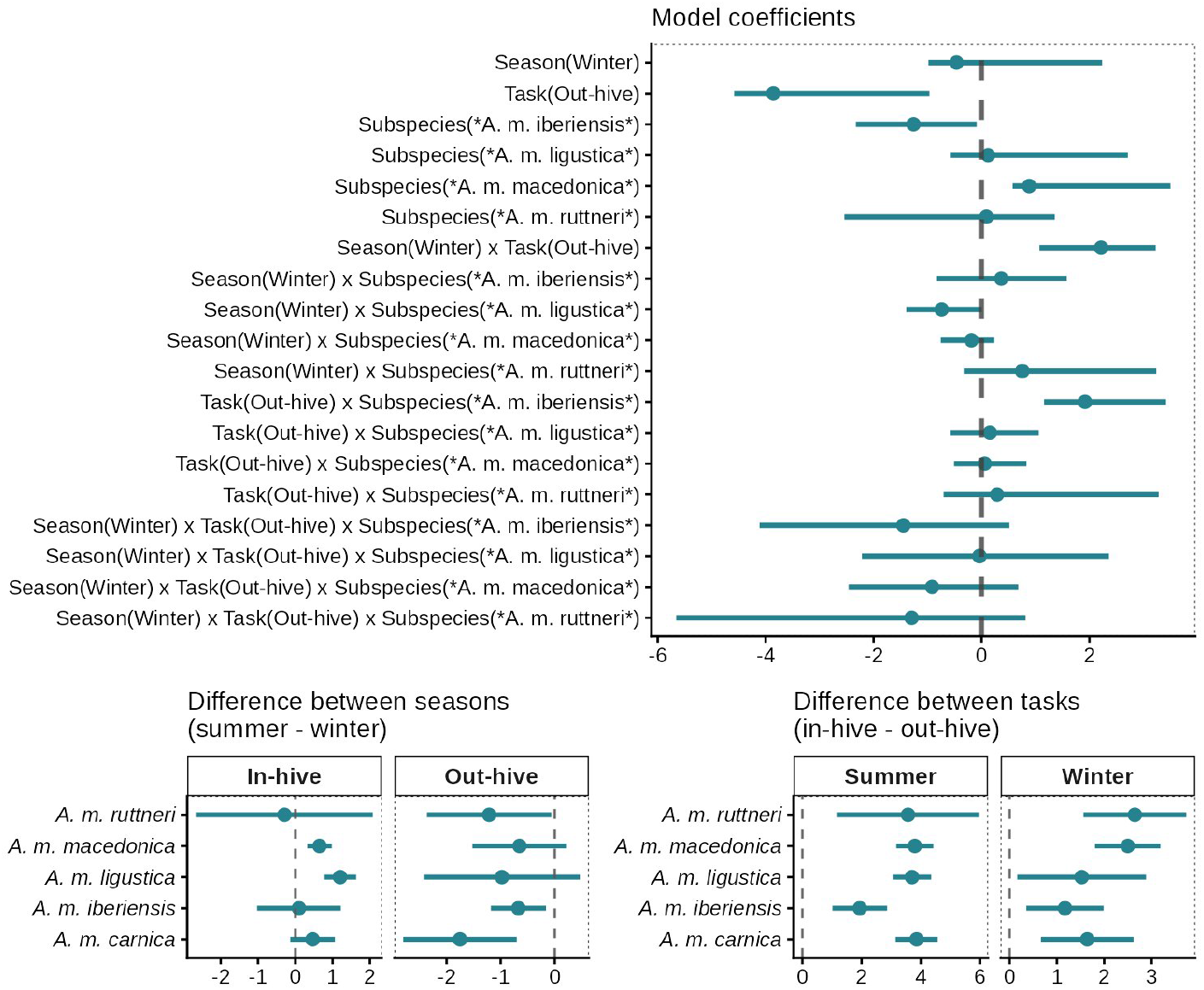
Mean hydrocarbon chain length differences in the CHC profiles of summer and winter in- and out-hive workers of five *A. mellifera* subspecies. The figure is divided into three plots depicting the quantile regression model coefficients (top), and the pair-wise contrasts between seasons (bottom left) and tasks (bottom right). Point intervals represent the model estimates and their 95% confidence intervals. The null values are visualized with vertical dashed grey lines.

